# Intrinsic OASL expression licenses interferon induction during influenza A virus infection

**DOI:** 10.1101/2025.03.14.643375

**Authors:** Joel Rivera-Cardona, Tarun Mahajan, Neeharika R. Kakuturu, Qi Wen Teo, Joseph Lederer, Elizabeth A. Thayer, Elizabeth F. Rowland, Kyle Heimburger, Jiayi Sun, Cera A. McDonald, Clayton K. Mickelson, Ryan A. Langlois, Nicholas C. Wu, Olgica Milenkovic, Sergei Maslov, Christopher B. Brooke

## Abstract

Effective control of viral infection requires rapid induction of the innate immune response, especially the type I and type III interferon (IFN) systems. Despite the critical role of IFN induction in host defense, numerous studies have established that most cells fail to produce IFNs in response to viral stimuli. The specific factors that govern cellular heterogeneity in IFN induction potential during infection are not understood. To identify specific host factors that license some cells but not others to mount an IFN response to viral infection, we developed an approach for analyzing temporal scRNA-seq data of influenza A virus (IAV)-infected cells. This approach identified the expression of several interferon stimulated genes (ISGs) within pre-infection cells as correlates of IFN induction potential of those cells, post-infection. Validation experiments confirmed that intrinsic expression of the ISG OASL is essential for robust IFNL induction during IAV infection. Altogether, our findings reveal an important role for IFN-independent, intrinsic expression of ISGs in promoting IFN induction and provide new insights into the mechanisms that regulate cell-to-cell heterogeneity in innate immune activation.

## Introduction

Successful control of viral infection relies on the effective activation of both innate and adaptive immunity. The type I/III interferon (IFN) system forms a central component of the innate antiviral immune response and is critical for restricting early virus dissemination^1–4^. During influenza A virus (IAV) infection, the induction of type I and III IFNs is primarily driven by detection of viral RNA through the dsRNA sensor retinoic acid-inducible gene I (RIG-I)^5–9^. Once secreted, IFNs engage their respective type I or type III specific receptors in both paracrine and autocrine fashions, initiating JAK/STAT-dependent signaling cascades that lead to the induction of hundreds of interferon-stimulated genes (ISGs)^2,10^, which collectively can inhibit various aspects of the viral life cycle^10–12^.

The induction of both type I and type III IFNs during influenza A virus (IAV) infection has been extensively described^3,9,13–16^. This process involves detection of viral RNA species by RIG-I, which interacts with the scaffolding protein MAVS. This interaction ultimately activates transcription factors, such as interferon regulatory factors 3 and 7 (IRF3, IRF7), leading to the synthesis and secretion of IFNs^6,15,17–19^. Despite the near-universal expression of the key components of this pathway, it is clear that a large majority of infected cells fail to express type I or type III IFNs^20–28^. Similar results have been observed in cells stimulated with synthetic immune agonists in the absence of virus-mediated immune antagonism^23,25,29^. These findings suggest that the inability to induce IFNs during infection is not only due to viral antagonism but also involves heterogeneity in the expression and/or function of intrinsic cellular factors required for IFN production.

To identify the host factors that underly cell-to-cell heterogeneity in IFN induction potential, we developed a novel single-cell RNA sequencing-based pseudotime analysis approach. This analysis allowed us to identify genes whose pre-infection expression levels within individual cells correlated with the post-infection IFN induction potential of those cells. This method revealed a correlation between the intrinsic, pre-infection expression of a subset of ISGs within single cells and the IFN induction potential of those cells following infection. We further show that low-level intrinsic expression of the ISG OASL is required for maximal IFN induction during IAV infection. Altogether, these data reveal that heterogeneity in intrinsic ISG expression regulates single cell patterns of IFN induction during the earliest stages of viral infection.

## Results

### Longitudinal scRNA-seq analysis of cellular gene expression during IAV infection

We and others have demonstrated that only a small subset of cells infected with IAV actually express type I and/or III IFN^20–25^. While some of this heterogeneity in IFN expression potential may be due to virus mediated antagonism of the IFN system^30^, similarly low IFN induction frequencies have also been observed following stimulation with chemical immune agonists^23,25,29^. These findings suggest that the failure of many cells to induce IFN may not be solely the result of virus-mediated antagonism but may also reflect intrinsic cellular heterogeneity in IFN induction potential.

We hypothesized that cellular heterogeneity in pre-infection state may result in only a subset of cells being “licensed” to turn on IFN in response to infection-associated stimuli. To explore this hypothesis, we examined how single cell transcriptional patterns changed over the course of IAV infection. We infected A549 cells at a multiplicity of infection (MOI) of 0.5 NP-expressing units (NPEU)/cell^31^ with the human seasonal H3N2 isolate A/Perth/16/2009 (Perth09). Following establishment of infection, we added NH_4_Cl to cell culture media to block secondary spread to ensure all infections were synchronized^32^. At different times post-infection, we sorted infected cells based on HA surface expression (**Figure 1A**) and used sorted cells to generate single-cell RNA sequencing (scRNA-seq) libraries.

**Figure 1.**
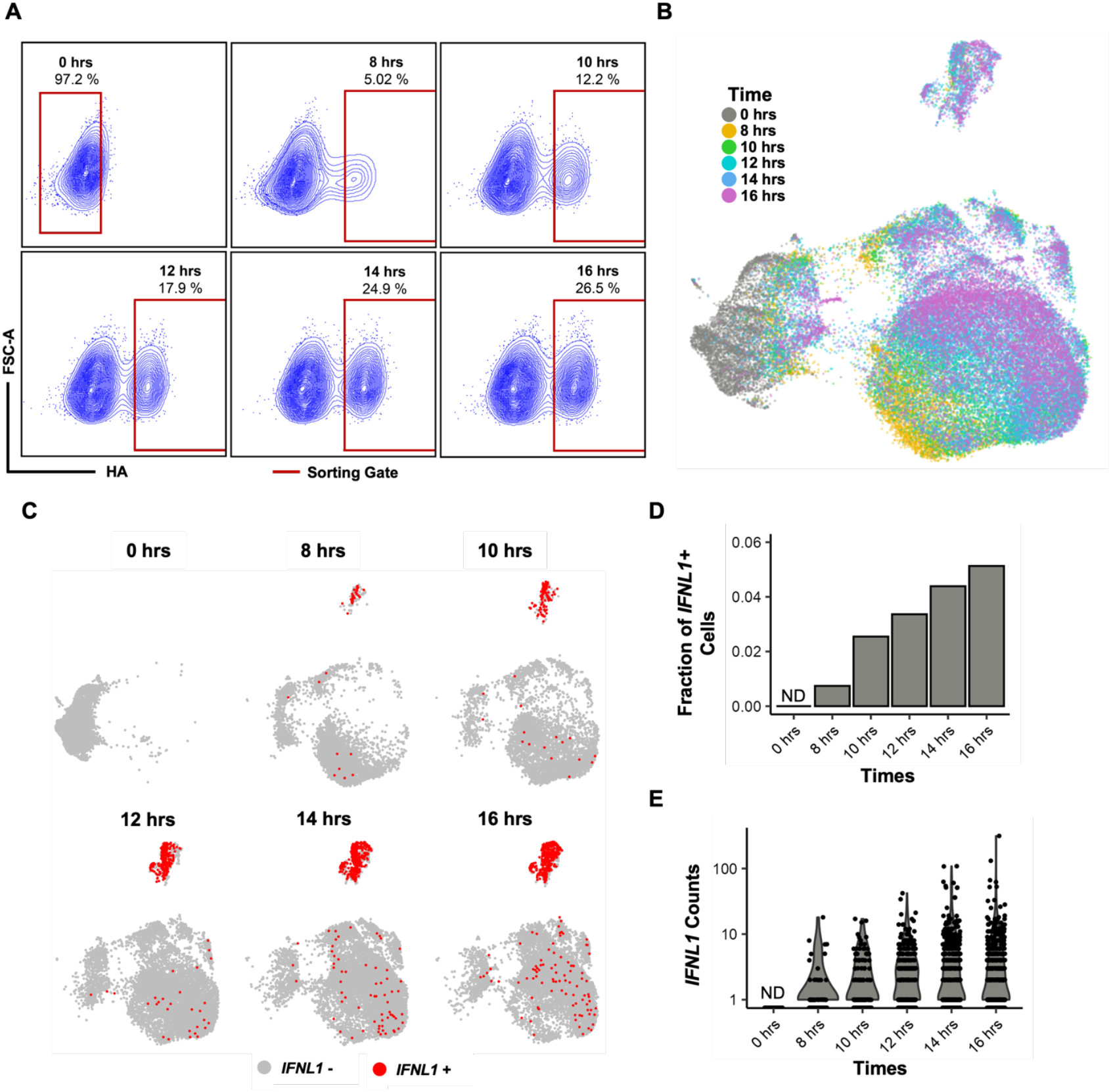
Temporal scRNA-seq analysis of A549 cells infected with H3N2. **A)** A549 cells were infected with Perth09 (MOI of 0.5 NPEU/cell) and collected at different times post-infection. Cells were sorted based on surface expression of the viral glycoprotein HA prior to library preparation for scRNA-seq. **B)** Collected scRNA-seq libraries were clustered based on transcriptional profiles and colored according to time. **C)** IFNL1-positive cells, highlighted in red, were identified at different times post-infection. **D)** Quantification of the fraction of infected cells expressing IFNL1 and **E)** expression counts at different timepoints.

We clustered cells based on transcriptional similarity and classified individual cells as infected or uninfected based on virus-mapping reads, as previously described^22,23^. We then identified cells with detectable *IFNL1* expression across all timepoints post-infection (**Figure 1B-C**). *IFNL1* transcript was detectable as early as 8 hrs post-infection in <1% of infected cells with maximum expression at 16 hrs with ∼5% of infected cells. The fraction of *IFNL1* positive infected cells and *IFNL1* transcript counts increased through the final 16 hrs timepoint (**Figure 1C-E**), similar to what we have previously reported during infection with this virus strain^23^.

Next, we examined the activation of type I and type III interferons, as well as interferon signaling mediated by autocrine and paracrine IFN sensing^2,33,34^. Type III interferons were expressed in a higher fraction of cells compared to type I interferons, potentially due to the early time point post-infection^3^ (**S1 Fig A, C**). Infected cells exhibited robust expression of interferon-stimulated genes (ISGs), which are primarily induced in response to IFN signaling (**S1 Fig B, D**). ISG expression was observed as early as 8 hrs post-infection and both the fraction of ISG-positive cells and expression levels increased over time.

### Pseudotime reconstruction of gene expression dynamics reveals divergent transcriptional trajectories during infection

To better understand what distinguishes the subset of infected cells that express IFN from the majority that do not, we generated a pseudotime-based reconstruction of the movement of infected cells through transcriptional state space. To do this, we adapted the RNA velocity method (noSpliceVelo^35^) to infer gene expression dynamics from scRNAseq data based on the temporal relationship between the expression mean and variance of individual genes. By applying noSpliceVelo to our longitudinal dataset, we could place individual cells in relation to each other within transcriptional state space and infer the transition of cells through this space over the course of infection. This approach enhanced our ability to calculate RNA velocity trajectories without relying on splicing dynamics, which may be disrupted during infection.

We next visualized the timepoints at which individual cells were collected and confirmed that our pseudotime reconstruction of gene expression trajectories accurately captured the expected transition from pre-infection transcriptional states to the distinct states observed at later timepoints (**Figure 2A**). To further evaluate whether our analysis recapitulated the presence of IFN+ and IFN-infected cells observed in previous studies^22,23^, we overlaid cellular transcriptional trajectories with *IFNL1* expression which highlighted trajectory of cells transitioning into IFNL positive or negative populations (**Figure 2B**). Our analysis revealed four distinct terminal transcriptional states, one in which type I/III IFN and other innate antiviral genes were highly expressed, and three where they were largely absent (**S2 Fig**). These findings confirmed that A549 cells diverge along two mutually exclusive transcriptional pathways following IAV infection, with only one leading to the induction of canonical innate antiviral genes such as interferons.

**Figure 2.**
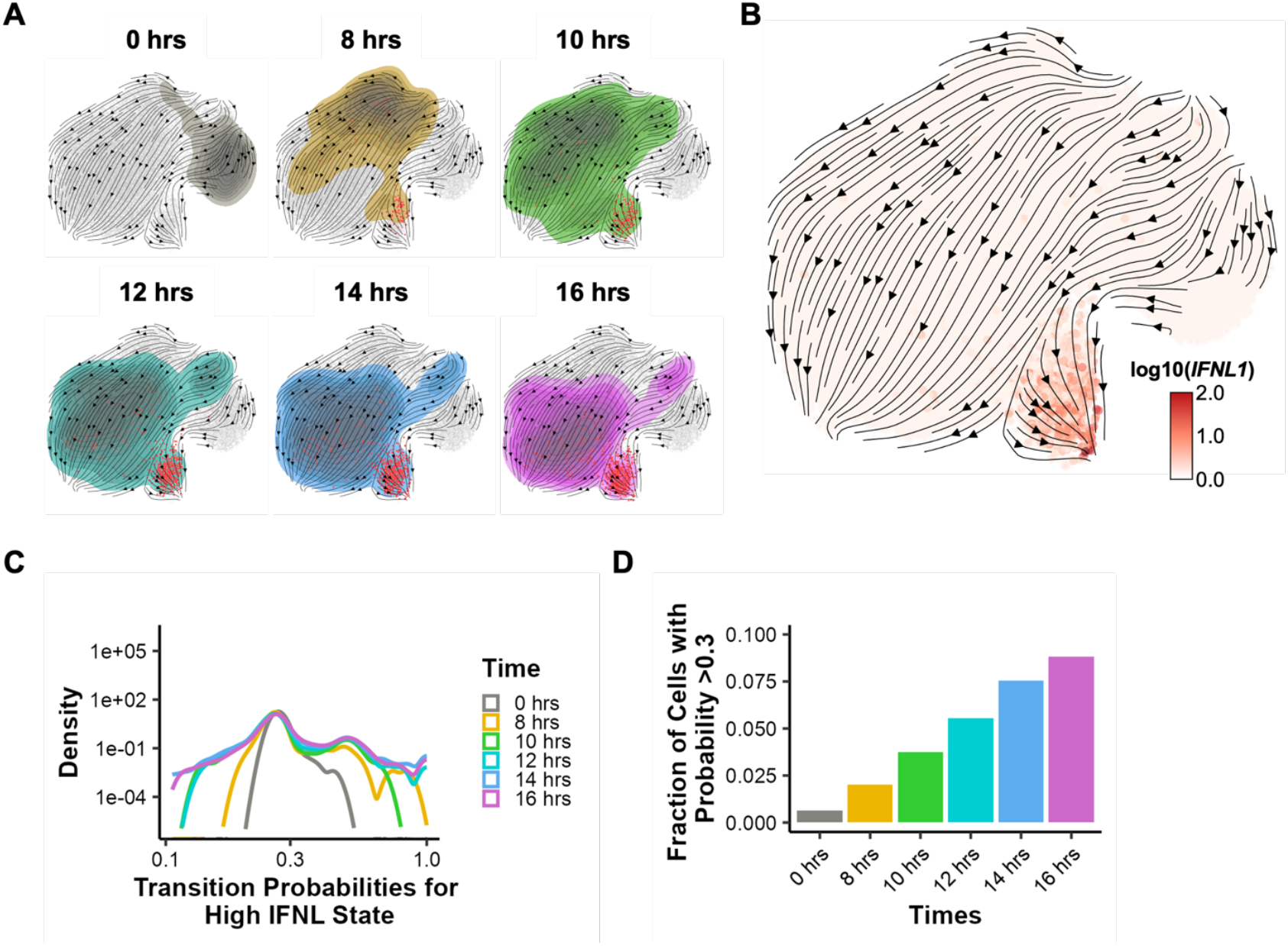
Identifying transcriptional terminal states and transition probabilities during H3N2 infection. **A)** Trajectory mapping of A549 cells transitioning through transcriptional state space, with cells from each timepoint represented by a distinct color and IFNL-positive cells highlighted in red. **B)** Trajectory map of A549 cells colored by IFNL1 expression log10(1+counts). **C)** Distribution of inferred probabilities for all cells transitioning into the high-IFNL terminal state. **D)** Fraction of cells at each timepoint with transition probabilities greater than 0.3 for reaching the high-IFNL terminal state.

This analysis approach allowed us to infer the probabilities that individual cells will transition into different terminal states (as previously described in CellRank2^36^). We estimated the probability of each cell in our dataset transitioning into the high IFNL state and plotted the probability distributions across the different timepoints (**Figure 2C**). As expected, these probability distributions broadened at later timepoints as individual cells approached their terminal states. We then determined the fractions of cells with transition probabilities higher than >0.3 for the high IFNL terminal state across timepoints (**Figure 2D**). Interestingly, we detected a small proportion of cells from the mock/0 hrs group with relatively high probability of transitioning to the high IFNL terminal state. These data suggest the presence of a subset of uninfected cells with elevated probability of expressing IFN during infection.

### Identification of pre-infection transcriptional correlates of IFN induction

We next assessed the correlation between individual cells’ probability of transitioning to the high IFNL terminal state and the expression levels of nearly 2,000 host genes. We identified 20 genes with the highest degree of correlation between expression level and transition probability in cells from the 0 hrs timepoint (**Figure 3A**). To examine potential functional relationships between these gene candidates, we performed a STRING database network analysis^37^, which highlighted three major clusters, one of which contained several well-known ISGs (**Figure 3B**). Several of these candidates play established roles in antiviral processes: *OASL* enhances RIG-I signaling^38^; *IRF1* maintains constitutive ISG expression, among other functions^39^; *ISG15* acts as a post-translational modification that regulates various antiviral pathways^40^; and *DDX60* promotes RIG-I binding to RNA^41^.

**Figure 3.**
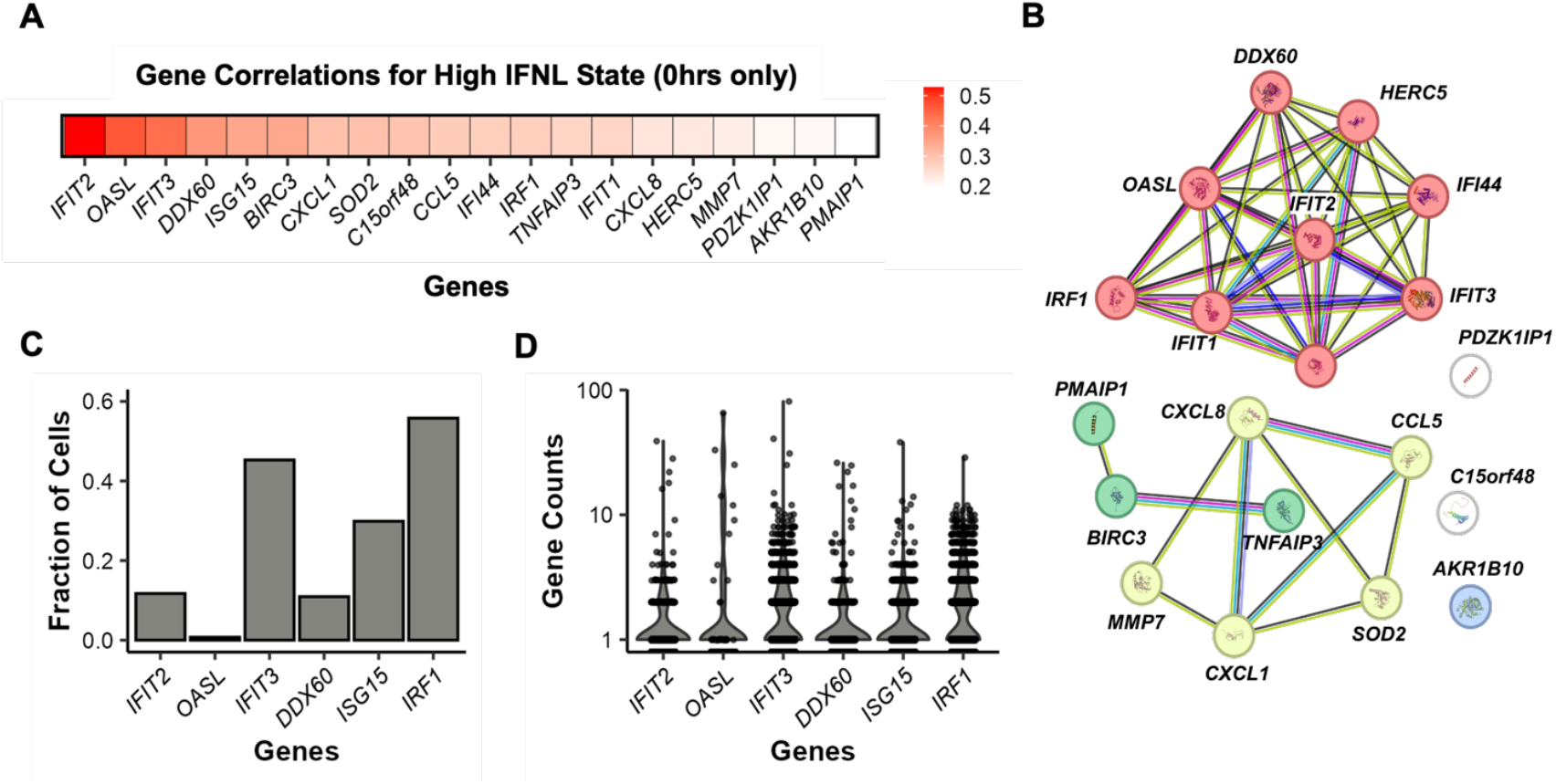
Identification of regulators of IFNL using pseudotime reconstruction analysis. **A)** Top 20 ranked genes according to correlation between inferred transition probabilities to the high IFNL terminal state and their expression at 0 hrs. **B)** STRING database network analysis of the top 20 correlated genes, colored by annotated cellular pathways. Genes in red cluster are defined as associated with interferon responses while the yellow and green clusters are associated with IL-10 and apoptosis respectively. **C)** Quantification of the fraction of cells expressing candidate genes at 0 hrs. **D)** Expression counts (log10) of candidate genes at the 0 hrs timepoint.

Next, to determine whether the candidate genes we identified were convincingly expressed in the absence of IFN signal or infection, we examined their baseline or intrinsic expression levels in our mock (0 hrs) sample. We observed substantial variability in expression frequencies for the candidate ISGs, with *IFIT3* and *IRF1* showing the highest percentages (∼40–50%) (**Figure 3C**). Each of these ISGs was highly expressed in a subset of uninfected cells despite the absence of IFN stimulation (**Figure 3D**). Altogether, these data suggested the existence of limited and highly variable intrinsic expression of some ISGs in subsets of uninfected cells in the absence of an active IFN response.

### Intrinsic *OASL* expression is required for early IFNL induction during IAV infection

Our top hit, *IFIT2*, has already been shown to enhance IAV replication^42^, so we did not pursue it further. Several of our other top hits (*e*.*g. OASL, IRF1, and DDX60)*, have previously been shown to modulate ongoing IFN responses, but have not been implicated in the induction of the IFN response^38,41,43–46^. For instance, following upregulation by IFN signaling, *OASL* can enhance RIG-I signaling, forming a feedback loop that potentiates ongoing IFN responses^38^. However, recent findings have observed low-level intrinsic expression of a subset of ISGs in the absence of IFN signaling, suggesting that some ISGs may serve functions in the absence or prior to IFN expression^44,45,47,48^.

To determine whether the ISGs that we identified as correlating with IFN induction potential play roles in modulating IFNL expression during IAV infection, we performed siRNA knockdowns of candidate genes for 48 hrs, followed by infection with Perth09 (H3N2) at an MOI of 1 NPEU/cell for 16 hrs. We measured secreted, biologically active, IFNL in the supernatant and found that knockdown of most candidate genes had no appreciable effect on IFNL secretion, with the exceptions of *OASL* and *IRF1*, both of which significantly reduced IFNL secretion and *DDX60* which resulted in increased IFNL secretion (**Figure 4A**).

**Figure 4.**
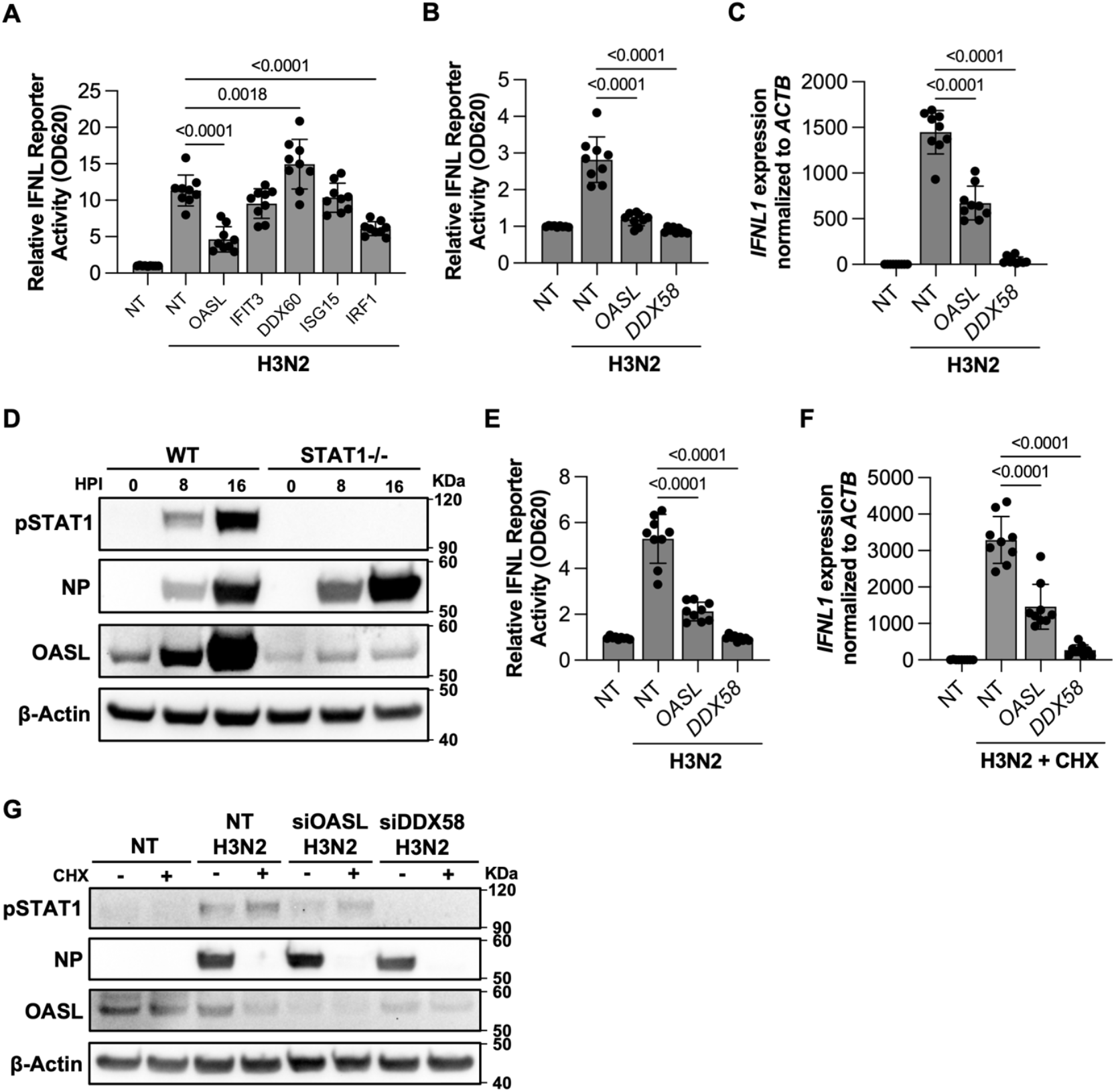
Intrinsic OASL is required for robust IFNL expression in response to IAV infection. **A)** Detection of IFNL in the supernatant of A549 cells transfected with siRNA targeting candidate genes for 48 hrs, followed by infection with H3N2 (MOI 1 NPEU/cell) for 16 hrs. **B)** Detection of active secreted IFNL and **C)** IFNL1 mRNA in siRNA-treated A549 cells infected at MOI 1 for 8 hpi. **D)** Detection of pSTAT1, NP, OASL, and β-Actin levels in A549 WT and STAT1-/-cells infected with H3N2 (MOI of 1 NPEU/cell) at 0, 8, and 16 hrs post-infection (hpi). **E)** Measurement of secreted functional IFNL in STAT1-/-A549 cells transfected with siRNA and infected at MOI 1 for 8 hpi. **F)** Quantification of IFNL1 expression in siRNA-treated A549 cells infected with H3N2 (MOI of 5 NPEU/cell) and subsequently incubated with cycloheximide (CHX) (10 µg/mL) for 8 hrs. **G)** Detection of levels of pSTAT1, NP, OASL, and β-Actin in A549 cells treated with CHX (10 µg/mL) or DMSO for 8 hrs. Data are shown as mean with SD; N = 9 (3 independent experiment with 3 replicates each). One-way ANOVAS with Dunnett’s multiple comparison tests were used for statistical analysis. Western blots are shown as a representative of three independent experiments.

While IFN signaling-dependent upregulation of OASL is known to help potentiate ongoing IFN responses by enhancing RIG-I activity^38,43^, OASL is not thought to play a significant role in the initial induction of the IFN response. To evaluate the importance of OASL in IFN induction in our system, we treated A549 cells with siRNA targeting *OASL, DDX58* (RIG-I) as a positive control, or a negative control siRNA for 48 hrs and then infected with H3N2 at an MOI of 1 NPEU/cell.

We measured secreted IFNL at 8 hpi, as we previously identified this as the earliest timepoint that IFNL could be detected during Perth09 infection^23^. We observed a significant reduction in IFNL levels in siOASL-treated cells compared to the non-targeting control (NT), with no detectable IFNL in siDDX58-treated cells (**Figure 4B**). Consistent with this, we measured a ∼3-fold reduction in *IFNL1* transcript levels in siOASL-treated cells compared with control siRNA treatment (**Figure 4C**).

The IFN-dependent upregulation of OASL has been previously shown to enhance RIG-I sensitivity and IFN induction^38^. It is possible that the effect of siRNA-mediated knockdown of OASL on IFN may simply be due to inhibiting OASL induction rather than reflecting any activity of intrinsically expressed OASL. To differentiate between these possibilities, we examined the effects of OASL knockdown in STAT1-/-cells, which are incapable of IFN signaling (**S3 Fig**). We infected WT and STAT1-/-A549 cells with H3N2 at an MOI of 1 NPEU/cell and measured levels of OASL and pSTAT1, a marker for activation of interferon signaling, by western blot at 8 and 16 hpi (**Figure 4D**). As expected, pSTAT1 was detectable by 8 hpi in WT but not STAT1-/-cells. OASL protein levels were elevated at 8 and 16hpi in WT cells, consistent with IFN-mediated induction; however, no induction of OASL was observed in the STAT1-/-cells. These data indicate that H3N2 infection can increase OASL expression by 8 hpi in a STAT1-dependent manner. Thus, the absence of functional JAK/STAT signaling allows us to specifically dissect the role of intrinsic OASL expression during infection.

We next used STAT1-/-cells to directly assess the role of intrinsic OASL expression in IFN induction. We treated STAT1-/-A549 cells with siRNA targeting *OASL, DDX58*, or non-targeting controls, infected with H3N2, and measured IFNL secretion at 8 hpi. Like wild-type (WT) A549 cells, siOASL-treated STAT1-/-cells exhibited a significant reduction in IFNL secretion compared to controls (**Figure 4E**), demonstrating that intrinsic OASL expression is critical for the early induction of IFN during IAV infection.

Finally, to completely rule out the possibility of virus-induced OASL contributing to IFN modulation, we examined the effect of OASL knockdown on IFN induction in the presence of cycloheximide (CHX), a potent translation inhibitor^49^. We treated A549 cells with siRNA targeting *OASL, DDX58* (RIG-I) as a positive control, or a negative control siRNA for 48 hrs before infection with H3N2 at an MOI of 5 NPEU/cell. Following infection, siRNA-treated A549 cells were incubated with CHX at a concentration sufficient to block detectable viral NP production (**S4 Fig**). In the presence of CHX, *IFNL1* induction in siOASL-treated cells was still significantly reduced compared to the negative control (**Figure 4F**). As expected, CHX treatment prevented the detectable accumulation of NP and prevented any increase in OASL levels above levels seen in uninfected control cells (**Figure 4G**). Together, these data demonstrate that intrinsic OASL levels at the time of infection are critical for IFNL induction. These findings further confirm the role of intrinsic OASL in modulating early IFN responses after stimuli.

### Intrinsic ISG expression within primary human respiratory tract epithelial cells

Previous studies have identified the intrinsic expression of various ISGs as a major factor contributing to the resistance of stem cells to viral infection^47^. However, little is known about the intrinsic expression of ISGs across differentiated cells within the respiratory tract or other tissue systems. To investigate the expression patterns of ISGs within untreated, uninfected human respiratory tract cells, we analyzed scRNA-seq datasets from primary human bronchial epithelial cells (HBECs) differentiated at the air-liquid interface. These HBEC cultures have been shown to serve as a physiologically relevant model for studying infection dynamics in the respiratory tract^50–56^. We examined the distribution of ISGs across different cell populations within four publicly available datasets of uninfected HBECs, two which have been previously used to study viral infection dynamics^52,56^.

We classified cell types within the HBEC cultures based on transcriptional profiles using ScType^57^ and identified five major populations present in all datasets: secretory cells (including goblet cells), basal cells, ciliated cells, club cells, and ionocytes (**Figure 5A and S5 Fig**). Additionally, we identified a unique population in three out of the four datasets, which we classified as cycling basal cells based on the population’s high expression of basal cell and proliferation-associated genes, including *KRT13, KRT5, KRT14, MKI67, TOP2A*, and *TP63*, consistent with previously reported markers for this cell type^58–61^ (**S6 Fig**).

**Figure 5.**
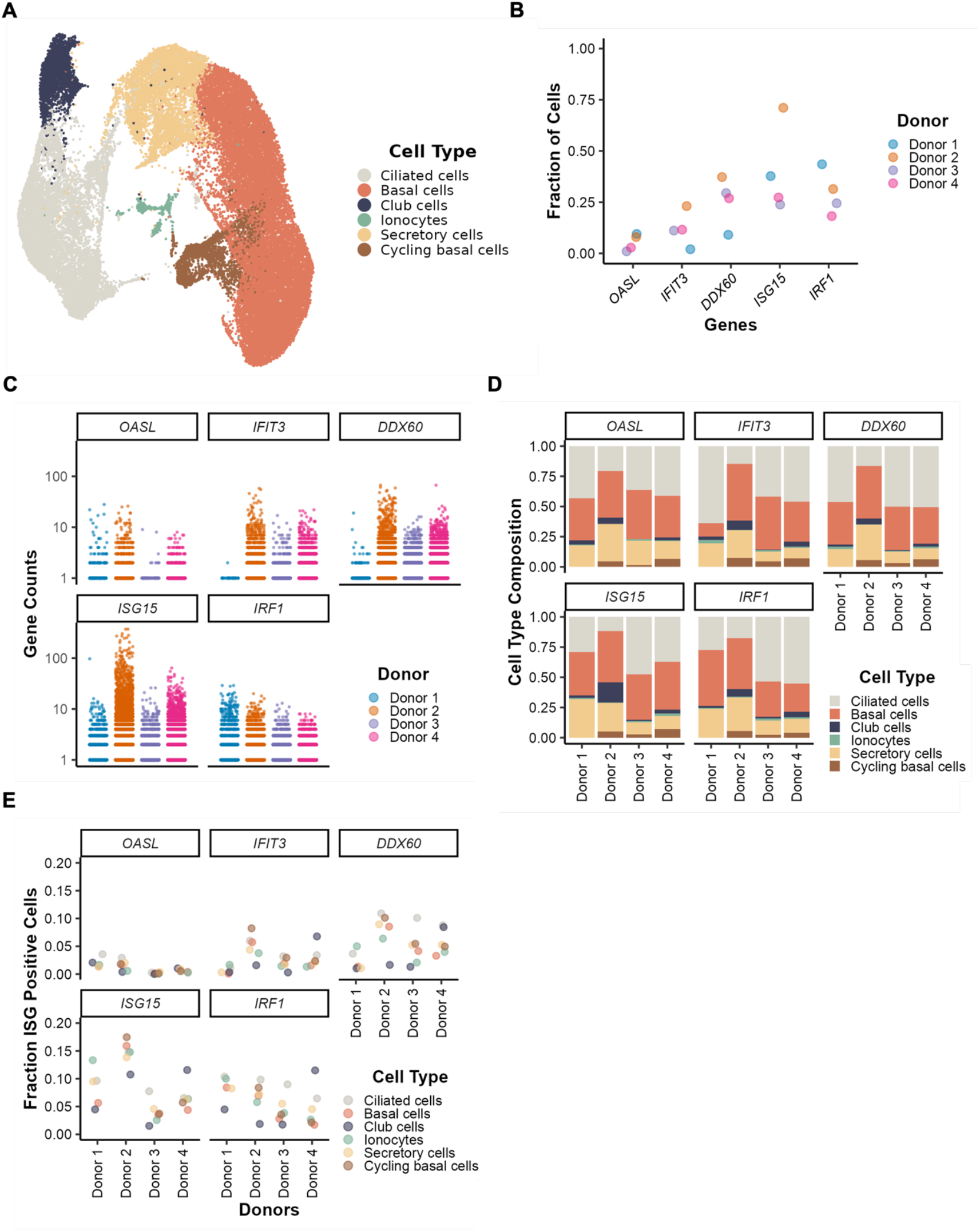
Identification of intrinsic ISGs in primary human bronchial epithelial cells by scRNA-seq. **A**) UMAP visualization of scRNA-seq data from HBECs, with cells labeled based on cell types. **B**) Proportion of cells expressing OASL, IFIT3, DDX60, ISG15, or IRF1 and **C)** expression counts divided by donor. **D)** The cell type composition of positive cells for each candidate gene separated and colored by cell types. **E)** Fraction of cells within each cell type expressing the candidate ISGs.

Next, we assessed intrinsic ISG expression levels across four donors, focusing on the ISGs identified in our pseudotime analysis (**Figure 5B-C**). Notably, a surprisingly high fraction of cells from diverse donors expressed these ISGs, albeit at relatively low levels. Expression patterns were largely consistent across donors, except for donor 2, who exhibited higher levels of *ISG15* compared to the others. Donor 1 had the lowest expression of *IFIT3* and *DDX60*; however, the library size was much smaller for this donor compared with the other three (**S5 Fig**), potentially skewing the cell frequency measurements for this donor. Interestingly, we observed substantial variation in *OASL, IFIT3, DDX60*, and *ISG15* expression, suggesting that some individuals may have inherently higher levels of intrinsic ISG expression, potentially influencing antiviral responses.

To further explore variability in intrinsic ISG expression, we examined the cell type composition of ISG positive cells across all four donors (**Figure 5D**). Ciliated, basal, and secretory cells were the primary cell types with high ISG expression. We further examined the fractions of each cell type that were positive for the individual ISGs (**Figure 5E**). While the donors generally exhibited similar proportions of cell types expressing the individual ISGs, there were some clear signs of significant donor-to-donor variability in intrinsic ISG expression. For instance, the ISG expression frequencies were highest in club cells in donor 4, while club cells generally exhibited the lowest ISG expression frequencies for the other three donors. Altogether, our results indicate that several ISGs, including *OASL*, exhibit heterogeneous intrinsic expression patterns within multiple human respiratory tract cell types, where they may act to regulate IFN induction during viral infection.

## Discussion

The apparent discrepancy between the critical importance of the IFN system for host defense and the surprisingly small percentages of infected cells that produce IFN has never been adequately explained. In this study, we developed a novel pseudotime reconstruction method to analyze longitudinal scRNA-seq data and used it to identify OASL as a key intrinsically expressed ISG that licenses RIG-I-mediated *IFNL1* induction within a subset of cells during IAV infection. Our findings demonstrate that heterogenous intrinsic ISG expression influences the IFN induction potential of individual cells, revealing an underappreciated layer of regulation that governs cellular heterogeneity in innate immune function.

IFN induction during IAV infection primarily occurs through RIG-I^15,62^. Maximal RIG-I signaling activity typically requires K63-linked polyubiquitination of its CARD domains^63–68^, but OASL can circumvent this requirement by facilitating RIG-I oligomerization and enhancing signaling activity^38,43,69^. Previously, this function had only been thought to operate in the context of an ongoing IFN response, following IFN-dependent induction of OASL. We hypothesize that intrinsic OASL expression increases baseline RIG-I sensitivity, allowing more efficient detection of IAV RNA during the early stages of infection before the virus is able to effectively antagonize IFN induction^30^. In this way, intrinsic OASL expression can be seen as licensing a subset of cells within a population to serve as IFN-competent sentinels during viral infection. In this model, the frequency of these responder cells within tissue would determine the extent of viral replication and spread that is required to trigger an IFN response. Our study focused on IAV-RIG-I interactions and IFN induction, but future work should explore the mechanisms governing cellular heterogeneity in responsiveness to other infection-associated stimuli and immune signals.

Our data expand what is known about how ISGs can function in antiviral defense. Intrinsic ISG expression has previously been shown to protect stem cells from viral infection^47^. We show here that intrinsic ISG expression also plays an important role in innate immune function in differentiated epithelial cells. What mechanism(s) govern the intrinsic expression patterns of OASL and other ISGs across cell populations at steady state? This remains an open but important question as we expect that the frequency of responder cells will influence susceptibility to both viral infection and tendency to develop IFN-driven autoimmune conditions. While our sample size is small, our data from primary human airway cells suggests that there may be substantial person-to-person and cell type variation in intrinsic ISG expression, and by extension, IFN induction potential. Ciliated cells are the primary targets for respiratory viruses such as influenza and SARS-CoV-2 as well as drivers of IFN expression^48,52,56,70,71^, potentially explaining the relatively high levels of intrinsic ISG expression we observed in these cells. The relatively high intrinsic ISG levels we observed within basal and cycling basal cells could potentially be related to their less-differentiated state^72^, as stem cells have previously been shown to express high levels of intrinsic ISGs^47^. Future studies should aim to define the specific regulatory features associated with intrinsic ISG expression and explore how these features vary for different ISGs across cell types, tissues, and individuals.

While we focused in this study on the role of OASL during IAV infection, our results raise important questions about how intrinsic OASL expression may function to regulate inflammatory processes in the absence of viral infection. Gain-of-function mutations that lower RIG-like receptor activation thresholds have been described to lead to more frequent recognition of self-RNAs and are associated with interferonopathies like Aicardi–Goutières syndrome^73–75^. Deficiencies in other key ISGs including ISG15, MX1, OAS/RNase-L have also been linked to the development of autoimmune disorders^73,76–80^. Based on the function of OASL we describe here it is easy to imagine that individuals with elevated frequencies of intrinsic OASL expression will be more likely to trigger inappropriate IFN responses to self-RNAs, thus increasing risk of developing a variety of interferonopathies. More work is clearly needed to better characterize the regulation and function of intrinsic ISGs like OASL in balancing host defense and the potential for immunopathology across diverse cell types and tissues.

## Material and methods

### Plasmids and cells

Reverse genetic plasmids for A/Perth/16/2009 were generously provided by Dr. Seema Lakdawala. Human lung epithelial cells (A549) and Madin-Darby canine kidney (MDCK) cells were obtained from Dr. Jonathan Yewdell and maintained in Gibco’s Dulbecco’s Modified Eagle Medium (DMEM) high glucose supplemented with GlutaMax and sodium pyruvate, and Gibco’s Minimal Essential Medium (MEM) with GlutaMax (Life Technologies), respectively. Human embryonic kidney (HEK293T) cells were provided by Dr. Joanna Shisler and maintained in Gibco’s MEM with GlutaMax. All growth media were supplemented with 8.3% fetal bovine serum (Avantor), and cells were cultured at 37°C with 5% CO_2_.

### Virus rescue and quantification

Recombinant A/Perth/16/2009 was generated using reverse genetics by transfecting HEK293T cells at 60–80% confluency with 500 ng of each plasmid encoding the viral genome segments, using jetPRIME (Polyplus). MDCK cells were added 24 hrs post-transfection in Gibco’s Minimal Essential Medium (MEM) with GlutaMax, supplemented with 1 mM HEPES (Life Technologies) and 1 μg/mL tosylsulfonyl phenylalanyl chloromethyl ketone (TPCK) trypsin (Worthington Biochemical Corporation, Lakewood, NJ). Supernatants were collected 48 hrs post-transfection, and viral titers were determined using the tissue culture infectious dose 50 (TCID_50_) assay. Working stocks were generated by infecting MDCK cells in T-75 flasks at a multiplicity of infection (MOI) of 0.001 TCID_50_/cell for 48 hrs, followed by viral titer quantification based on NPEU (NP-Expressing Units). NPEU/mL for virus stocks were calculated as previously described^31^, by infecting A549 cells with serial dilutions of virus under single cycle conditions and then measuring the numbers of cells expressing viral nucleoprotein (NP) using the anti-NP mouse monoclonal antibody HB65 by flow cytometry (BD FACSymphony A1).

### Viral infections

A549 cells were infected at an MOI determined by NPEU/cell by diluting the virus in PBS+/+ with 0.1% BSA in 24-well plates and incubating for 1 hrs at 37°C. Following incubation, the virus was removed, and cells were washed with PBS-/-. Cells were then cultured in Gibco’s DMEM (high glucose, HEPES, no phenol red; Life Technologies) supplemented with 8.3% fetal bovine serum and 6.5 μg/mL C05 anti-H3 neutralizing antibody to prevent secondary spread.

For scRNA-seq library preparation, A549 cells were infected with A/Perth/16/2009 at an MOI of 0.5 NPEU/cell. The virus was diluted in PBS+/+ with 0.1% BSA and incubated in 6-well plates for 1 hrs at 37°C. After incubation, the virus was removed, and growth media was added. At 3 hrs post-infection, the media was replaced with F-12 (Life Technologies) supplemented with 50 mM HEPES and 20 mM NH_4_Cl to inhibit secondary spread. Cells were collected at 8, 10, 12, 14, and 16 hrs post-infection, stained with the HA stem-specific antibody FI6v3, and sorted using a Bigfoot cell sorter.

### Single cell RNA-seq analysis

A549 cells were sorted, and viability was assessed using a BD20 cell counter (Bio-Rad). Cells were then diluted to 5,000 cells per sample, and libraries were constructed using the Chromium Next GEM Single Cell 3′ Reagent Kits v3.1 (Dual Index) (10x Genomics). Samples were sequenced on a NovaSeq 6000 (Illumina) using one S4 flow cell with a 2×150 nt read configuration.

HBECs were dissociated into single cell suspensions. Cell count and viability was assessed with the LUNA-FL cell counter (Logos Biosystems). A target of 10,000 live cells were captured per sample for Donors 1 and 2, 15,000 for Donor 3, and 20,000 for Donor 4. GEM generation, indexing, and library preparation was done using 10X Chromium Single Cell 3’ Reagent Kits. Sequencing for Donors 1 & 2 was run on NovaSeq S2. at 2×100 nt PE reads. Donors 3 & 4 were run on NovaSeq X (Illumina) and NovaSeq X Plus (Illumina) respectively, both using 2×150 nt PE reads.

### siRNA treatment and quantification of gene expression

Pooled siRNAs were purchased as ON-TARGETplus SMARTpool (Dharmacon Reagents) and reconstituted in nuclease-free water (Ambion). Knockdown efficiency was assessed by transfecting A549 cells with Lipofectamine RNAiMAX (Invitrogen) for 48 hrs, following the manufacturer’s instructions. After incubation, RNA was extracted using the RNeasy Mini Kit (Qiagen), and cDNA was synthesized using the Verso cDNA Synthesis Kit (Thermo Scientific). Reactions were setup by combining 4 μL of 5x cDNA synthesis buffer, 2 μL of dNTPs, 1 μL Oligo-dT, 1 μL of RT enhancer, 1 μL of Verso enzyme, 5 μL of cDNA and 6 μL of nuclease-free water. Reactions were incubated at 45°C for 50 min and 95°C for 2 min and held at 4°C; cDNA was stored at -20 and subsequently used for quantitative PCR. qPCR reactions were setup by combining 10 μL of TaqMan fast advanced master mix (Applied Biosystems), 1 μL of probe (*OASL;* Applied Biosystems: Hs00984387_m1, *IFIT3*; Applied Biosystems: Hs01922752_s1, *DDX60*; Applied Biosystems: Hs01102712_m1, *ISG15*; Applied Biosystems: Hs01921425_s1, *IRF1*; Applied Biosystems: Hs00971965_m1), 1 μL of cDNA and 8 μL of nuclease-free water. Cycle conditions were set as follows 50°C for 2 min, 95°C for 2 mins, and 40 cycles of 95°C for 1 second followed by 60°C for 20 seconds in a QuantStudio 3 (Applied Biosystems).

Knockdowns were performed by treating A549 cells with 15 μM siRNA per well (unless otherwise specified), in 24-well plates for 48 hrs. Following incubation, cells were infected as described previously, and RNA was extracted for cDNA synthesis. Gene expression of *IFNL1* (Applied Biosystems: Hs00601677_g1) was quantified by qPCR and normalized to *ACTB* (Applied Biosystems: Hs01060665_g1).

### Detection of secreted IFNL

Supernatants from infected A549 cells were collected and incubated with HEK-Blue IFN-λ reporter cells (InvivoGen) in 96-well plates for 24 hrs. Following incubation, the supernatant from HEK-Blue cells was clarified by centrifugation at 2,000 rpm for 5 minutes at 4°C. A volume of 20–50 μL was then incubated with QUANTI-Blue reagent (InvivoGen) according to the manufacturer’s protocol. Optical density at 620 nm was measured every 30 minutes for up to 2 hrs.

### CRISPR knockout of A549 cells

STAT1 knockout A549 cells were generated using single-guide RNA (sgRNA) 5’-CACCGAGAACACGAGACCAATGGTG-3’ and 5’-AAACCACCATTGGTCTCGTGTTCTC-3’. These were cloned into lentiCRISPR v2 (Addgene # 52961) as previously described^81^, followed by transfection into HEK293T cells using JetPRIME. Lentivirus was harvested 72 hrs post-transfection and A549s were infected in the presence of polybrene (Sigma-Aldrich). After 48 hrs virus was removed, and cells were selected in 1 μg/mL puromycin (InvivoGen). Bulk A549 population was validated by Western blot and maintained in 0.5 μg/mL puromycin.

### Western blot analysis

Cell lysates were collected by incubating A549 cells with RIPA lysis buffer supplemented with a protease and phosphatase inhibitor cocktail (Thermo Scientific) for 15 minutes at 4°C. Following incubation, samples were centrifuged at 20,000×g for 15 minutes. Protein concentrations were determined using a BCA microplate assay (Thermo Scientific) according to the manufacturer’s instructions.

For Western blot analysis, 10–20 μg of protein per well was loaded and transferred onto a PVDF membrane using the iBlot 2 dry transfer system (Invitrogen). Membranes were blocked with StartingBlock blocking solution (Thermo Scientific) and incubated overnight with primary antibodies: Rabbit anti-β-Actin (4970; Cell Signaling), Rabbit anti-OASLp (36845; Cell Signaling), Rabbit anti-pSTAT1 (9167; Cell Signaling), and Rabbit anti-NP (GTX125989; GeneTex). After incubation, membranes were washed with PBS + 0.1% Tween and incubated with an HRP-conjugated secondary antibody (Goat anti-Rabbit HRP; G-21234; Invitrogen) for 1 hrs at room temperature and visualized using the iBright 1500 imaging system.

### Human bronchial epithelial cells (HBECs)

HBECs scRNA-seq libraries originated from two published and two unpublished studies and were obtained from NCBI GEO. Cells were differentiated from four different healthy donors; Donor 1: Hispanic male 56 years, Donor 2: Black female 19 years, Donor 3: Hispanic male 43 years, and Donor 4: Caucasian male 55 years. Donor information was generously provided by Dr. Ryan A. Langlois.

HBECs (Lonza Bioscience) were harvested from healthy individuals. Cells were expanded using Pneumacult-Ex Plus Media (STEMCELL) and seeded onto 6.5mm transwell 0.4μm pore polyester inserts (STEMCELL). The transwell cultures were airlifted once confluent: apical media removed to expose cell culture surfaces to air and basal compartment medium replaced with Pneumacult-ALI Basal Media (STEMCELL). The cultures were maintained at air-liquid interface (ALI) for 3 weeks in a humidified incubator at 37°C and 5% CO_2_ before use.

### Biophysical model of bursty gene expression

The biophysical model central to noSpliceVelo^35^ and the analysis implemented in this study has been previously experimentally validated^82–87^. In this model, mRNA is transcribed in bursts, whose length are geometrically distributed with an average burst size equal to *β*. Next, the parameter *f* is used to denote the burst frequency. Each mRNA molecule is assumed to be degraded by a first-order process with rate *γ*. Biophysically, the burst length *β* and burst frequency *f* determine the kinetics of gene expression. The product of f and *β* determines the expression rates [1]. This bursty model can be solved to obtain differential equations for the time dependence of *σ2(t)* and *µ(t)*^35^, which we have recapitulated in eq. (1)

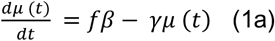

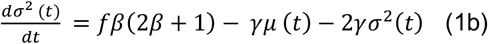

We previously showed that in these differential equations *σ*^2^(*t*) increases faster than *µ* by a factor of 2*β* + 1. For bursty genes (*β* ≫1), this allows the separation of up and down-regulated gene which can be used to inferred RNA velocity^35^.

### Selection of monotonic genes

NoSpliceVelo infers RNA velocity by leveraging genes which are up- or down-regulated and disregards the other genes. Separating up-from downregulated genes enabled dynamical modeling of bursty gene expression. Therefore, as the first step, we identify genes with monotonic kinetics over time. For this, we estimated *σ*^*2*^ and *µ* on a cell-by-gene level from scRNA-seq count data using a variational autoencoder (VAE)^35^. Briefly, for every cell, the encoder of this VAE takes as input the vector of raw gene counts and encodes it into a low-dimensional (10 dimensions) latent space; ten is the suggested dimension for the latent space for VAE-based estimation from scRNA-seq data^88,89^. The decoder then takes this latent vector and decodes it into the parameters of negative binomial distribution for each gene. The model is trained in a self-supervised fashion to maximize the likelihood of observed mRNA counts. We extracted the cell-by-gene mean and variance parameters for the negative binomial distributions from the trained model and used them as approximations for *µ* and *σ*^*2*^. The VAE used here is an extension of one previously characterized in scVI^88^.

For bursty genes the curvature of the *σ*^2^-vs-*µ* phase plot is pronounced and thus we filtered genes by burst size *β*. However, since we have not yet incorporated dynamics in the model, we estimated *β*_ss_ by assuming that the cells are at steady state using *µ* and *σ*^2^ via the following relationships:

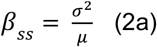

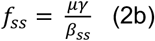

For each gene, we then took the mean of *β*_ss_ across cells and only retained those genes which have high burst rates (e.g., average *β*_ss_ ≥ 3). After this filtering, we were left with 1783 genes.

Next, to select monotonic genes, we focused on those genes which have at least weakly correlated (Pearson’s correlation coefficient (PCC) ≥ −0.3) averages *β*_ss_ and *f*_ss_ across experimental time. Since we exclude genes with large negative correlations, for the selected genes, either *f*_ss_ or *β*_ss_ or both can be expected to either monotonically increase or decrease over time. This filtering led to 88 up- and 463 down-regulated genes. We used these 551 genes to train the analysis implemented in this study.

### Model assumption

For our analysis we assume that post-infection any given gene is spontaneously either up- or down-regulated in *β* and/or *f*. We further assume that the genes do not exhibit non-monotonic behavior in the *σ*^*2*^-vs-*µ* phase space and thus we impose this constraint by filtering for up- and down-regulated genes, as described previously.

### Neural network architecture

noSpliceVelo decodes a state assignment variable k from the latent space z, which assigns to every cell in each gene the probability of belonging to the up-regulated branch of gene expression^35^. We modified the architecture of noSpliceVelo by making the state assignment variable k a known input variable (estimated when selecting monotonic genes) and removing the fully connected neural network from the latent space z to *s*. Since we apply this analysis to only monotonic genes, we removed the variable for switching time from the noSpliceVelo architecture.

### Dynamical model

Since we have already separated up- and down-regulated genes using the experimental data sampling times, we may use a simpler model than noSpliceVelo. To this end, we removed the decoder network for separating up- and down-regulated trajectories from noSpliceVelo. For modeling the dynamics of each gene, we used either the decoder for up- or down-regulated kinetics but not both. In contrast, noSpliceVelo utilized both decoders for all the genes, and aggregated the reconstructed means and variances of gene expression weighted by the estimated probabilities for assignment to up- and down-regulated trajectories.

Another difference between noSpliceVelo and the approach used here is the availability of experimental time. In noSpliceVelo, the outputs of the time decoder for all cells were clamped at the same maximum time. However, for each cell the output of the time decoder is clamped at the observed experimental time. This strategy imposes the temporal order from the experimental time but still allows cells from the same experimental timepoint to have different time assignments. For instance, two cells from the same experimental time sample, which are in the up-regulated trajectory of gene expression, could be at different time points along the trajectory. Low resolution of the experimental time would bin these two cells in the same time cluster.

### Direct modeling of raw mRNA counts

Since for our analysis the assignment of genes to up- and down-regulated trajectories of *σ*^*2*^-vs-*µ* phase space is determined using the pseudo steady state estimates of *f*_ss_ and *β*_ss_ (in conjunction with the experimental time), it can directly incorporate raw mRNA counts. Consequently, we model cells as transitioning from one steady state to another. In the absence of experimental time, noSpliceVelo needs two variables (*σ*^*2*^ and *µ*) to separate up- and down-regulated genes during model training.

### Generative process

The generative process for this analysis is similar to that of noSpliceVelo except that it models raw mRNA counts while the latter model’s variance and mean of gene expression. For data generation, our method first samples a random 10-dimensional vector, z, from a unit normal distribution. This vector is then decoded into *β, f*, and time *t* via fully connected neural networks. Similar to noSpliceVelo, separate fully connected neural networks decode *β* and *f* for the initial and final steady states. Consequently, *β* and *f* are cell and gene specific. In response to infection, the transcriptional kinetic parameters *β* and *f* could be dynamically altered and also potentially exhibit cell-to-cell variability.

Positivity is ensured for the parameters with the use of softplus activation. We further impose the following constraints on *β* and *f* for up-regulated genes with state assignment k_g_ = 0 (which is known from the step of selection of monotonic genes) to ensure monotonicity

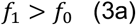

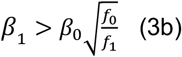

where the subscripts 0 and 1 represent the initial and the final steady states, respectively. For the down-regulated genes with state assignment *s*_*g*_ = 1, the direction of the constraints is flipped, and we get

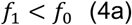

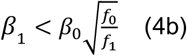

The time parameter *t* represents the time post-infection. To ensure that the inferred time is consistent with the experimental time, we used softplus activation after the fully-connected neural network for *t* and then clamped the output by the experimental time. This ensures that the predicted time follows the temporal ordering of the experimental time. Further, this still allows cells within the same experimental time cluster to differ in their dynamical time: experimental time has low resolution and hence, one time point measurement can reflect the properties of cells which represent different time points along the gene expression trajectory. The degradation rate *γ* is assumed to be only gene-specific and treated as a variational parameter.

Once we generate the time *t* and the kinetic parameters (*f* and *β*) at the initial and final steady states, for any cell, gene-specific *µ*(*t*) and *σ*^2^ can be generated using the time-dependent solution for *µ*(*t*) and *σ*^2^ for the bursty model of gene expression. These solutions have been reproduced from previous work^35^ in eq. (5) that follows:

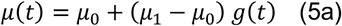

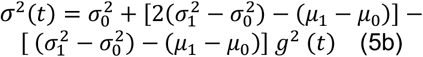

where *g* (*t*) = 1−e^−*γt*^ varies between 0 and 1 as time *t* goes from 0 to ∞. Also, *µ*_0_ and *µ*_1_ are the means of expression at the initial and the final steady-states, respectively. Similarly, 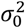 and 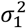 are the variances of expression. μ_0,_ μ_1_ and 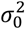 are obtained from *f*_0_, *f*_1_, *β*_0_ and *β*_1_ via eq. (2). After this point, the method diverges from noSpliceVelo, and samples mRNA counts rather than the mean and variance of expressions. With *µ*(*t*) and *σ*^2^(*t*) from eq. (5), mRNA counts are sampled from the negative binomial distribution.

### Inference procedure

The inference procedure for this analysis is similar to noSpliceVelo. Following the approach described in^89^, we use variational inference to determine (1) posterior probability distributions for the latent variables, (2) point estimates for the degradation rate, and (3) point estimates for the neural network parameters. After performing inference, velocity is computed as a functional of the variational posterior.

### Variational posterior

Unlike for noSpliceVelo, in our model the state assignment *s* is a known input variable, and the posterior distribution for *z* is given by

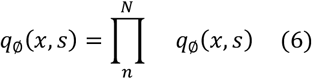

In eq. (6), the latent variable *z* factorizes over all cells and dependencies are captured via neural networks with parameters *ϕ*.

For likelihood computation, we use the negative binomial (NB) distribution to model raw mRNA counts^86^:

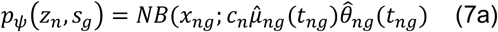

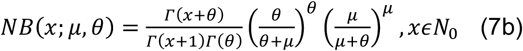

where 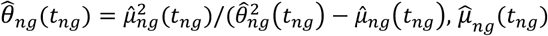, and 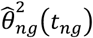 are given by eq. (5a) and eq. (5b), respectively. Furthermore, *f*_0_, *f*_1_, β_0_, and β_1_ are constrained by eqs. (3) and (4) for up-(*s*_*g*_ = 0) and down-regulated (*s*_*g*_ = 1) genes, respectively; *ψ* is the set of generative parameters, including the degradation rate γ and neural network parameters *ϕ*. In eq. (7a), c_n_ is the capture efficiency which was previously used for noSpliceVelo [1] and it is approximated using the 2000 least-variable genes (LVG) from the raw count matrix X using the expression given in^90^:

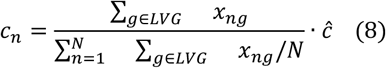

where ĉ is the average capture efficiency in the scRNA-seq protocol. We set *ĉ* = 0.1 for all our experiments.

### Loss function

For the training, the loss function is defined as

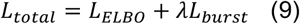

The negative of the evidence lower bound (ELBO)^91^ *L*_*ELBO*_ is given by:

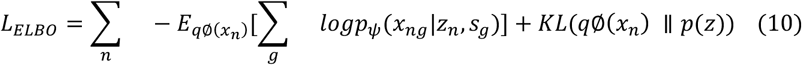

This can be estimated using randomly sampled minibatches of the data. The regularization term *L*_*burest*_ is designed to constrain the variability of burst size and frequency across different cells. To achieve this, we incorporate gene-specific variational parameters, namely *f*_1*gene*_, β_1*gene*_, *f*_0*gene*_ and β_0*gene*_. Rather than directly regulating the burst parameters, we ensure that *ℒ*_*burest*_ aligns with the scale of *L*_*ELBO*_by comparing the mean and variance, which are transformed using equations (2) a,b.

Finally, regularization is introduced by minimizing the discrepancy between the gene and cell-specific means (or variances) inferred from the latent cellular space and the corresponding gene-specific values. *L*_*burest*_ comprises four regularization terms:

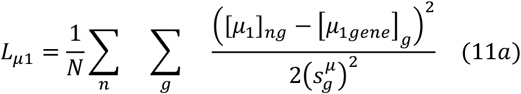

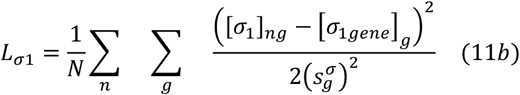

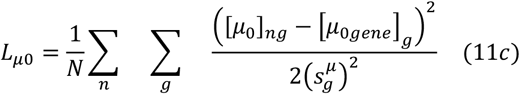

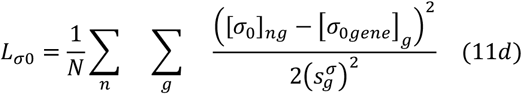

where terms with ‘gene’ (g) in the subscript are derived from gene-specific variational burst parameters via eqs. (2a,b), while 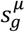 and 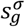 are variational parameters for adaptive scaling of the burst loss terms. Thus, *L*_*burest*_ = *L*_μ1_ + *L*_σ1,_+*L*_μ0_ + *L*_σ0_. In all experiments, we set *λ* = 10^−6^.

### Training

We configured the Training module following the same settings and default parameters outlined in^89^. The neural networks employed in this framework are fully connected, with ReLU activation in the hidden layers and softplus activation that ensures non-negative distributional parameters.

### Downstream tasks

Like in noSpliceVelo^35^ and other RNA velocity frameworks^89^, we used posterior predictive inference to estimate the mean and variance of expressions fitted by the model to infer RNA velocity. The steps for this are as follows:

1. Sample *z*_1_ from *q*_σ_ (*z*_1_ |*x*_1_)

2. Compute 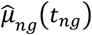 and 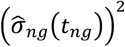 from eqs. (5a) and (5b), respectively, where the constraints for initial and final states are given by eq. (4a) or eq. (4b) if the gene is up- or down-regulated, respectively. Also, compute the velocity of the mean and variance of expression via equations (1a) and (1b) based on the inferred parameters.

3. Repeat steps 1 to 2 100 times and average the values of 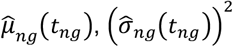, as well as velocities across these repetitions.

### Generating UMAP representations from the latent dimension

UMAP representation of the data was generated based on the latent dimension, using the Scanpy package^92^ in Python.

### Velocity streams

Velocity streams were generated by projecting the inferred RNA velocity onto the above described UMAP representation, using the method described in the splicing-based RNA velocity method scVelo^93^.

### Estimation of fate probabilities

To estimate probabilities of transition to different terminal states, we use the tool CellRank 2^36,94^. We use the VelocityKernel class with the mean velocity estimated, which we combined with the similarity-based ConnectivityKernel using the default relative weighting of 0.8 and 0.2, respectively. The state transition probability matrix between cells was estimated using this combined kernel. Next, as suggested in CellRank^36^, we use the Generalized Perron Cluster Cluster Analysis (GPCCA)^95^ to obtain macrostates, which may include potential initial and terminal cellular states. We determined terminal states based on visual inspection of the estimated macrostates. Finally, we computed the fate probabilities for every cell to transition to the assigned terminal cellular states.

## Data availability

The longitudinal A549 scRNA-seq dataset used in this study can be found in NCBI GEO with accession number GSE287024. Human bronchial epithelial cell scRNA-seq datasets were obtained from NCBI GEO GSE157526^52^, GSE247979^56^, and pending. In-house analysis code can be found in https://github.com/BROOKELAB/Temporal-scRNA-seq-H3N2.

## Acknowledgments

We are grateful to Dr. Jenny Drnevich from HPCBio and to the DNA Services Core at the Roy J. Carver Biotechnology Center for their support in preparing scRNA-seq libraries and conducting the initial data analysis. C.B.B was supported by NIH R01 AI139246, NIH R01 AI179910, and DOD DARPA INTERCEPT-W911NF-17-2-0034. R.A.L. was supported by NIH R01 AI148669. C.A.M. was supported by NIH T32 HL007741. O.M. would like to acknowledge funding from the NSF 1956384. The work of O.M. and S.M. was also supported in part by NSF 2107344. N.C.W was supported by NIH R01 AI165475.

## Supplementary figures

**S1 Fig.**
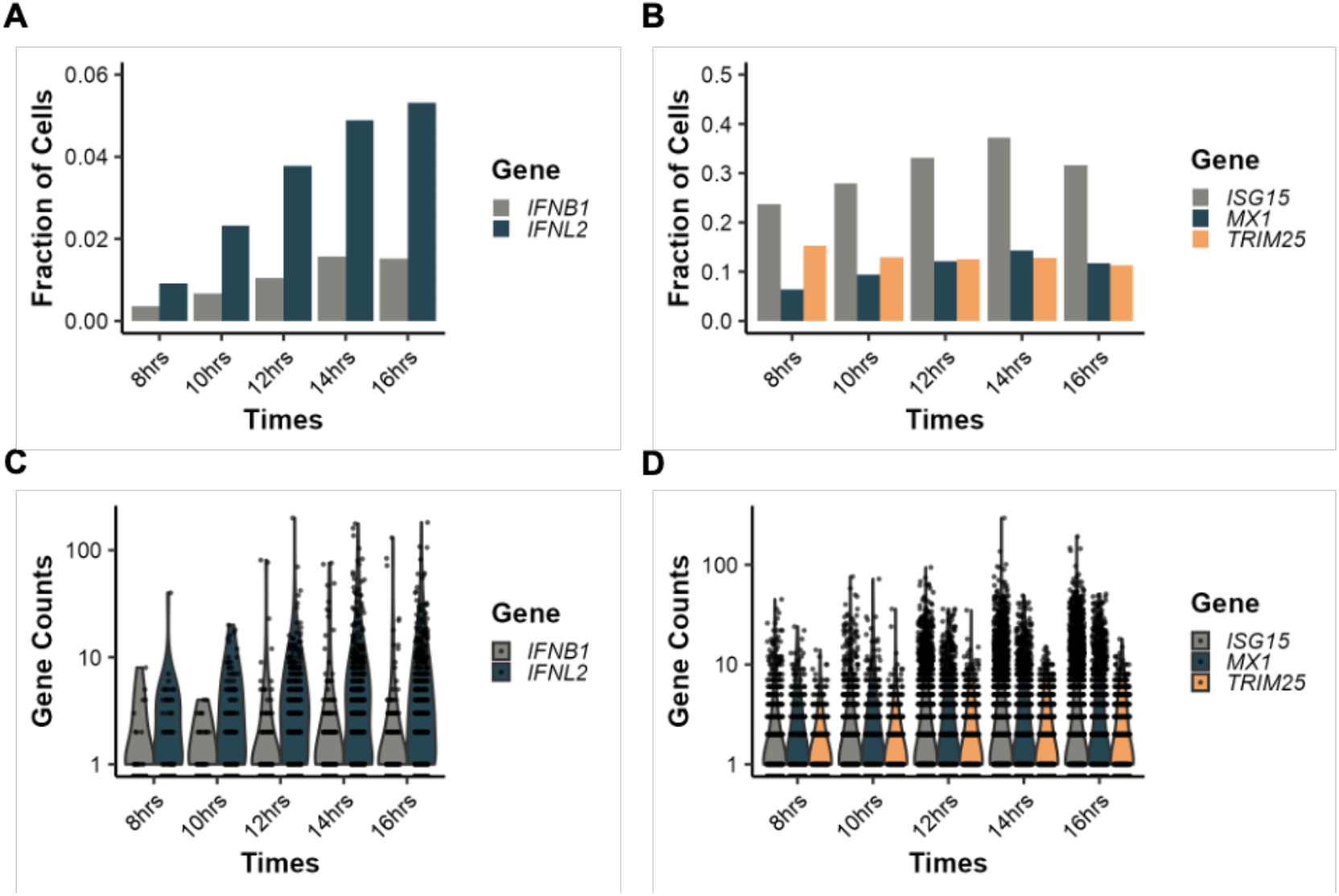
Quantification of diverse IFNs and ISGs in scRNA-seq libraries from H3N2-infected A549s. **A)** Fraction of infected A549s expressing type I and type III IFNs across different timepoints. **B)** Quantification of the fraction of infected cells expressing diverse ISGs at different times post-infection. **C)** Expression counts (log10) for type I and type III IFNs and **D)** ISGs in infected cells.

**S2 Fig.**
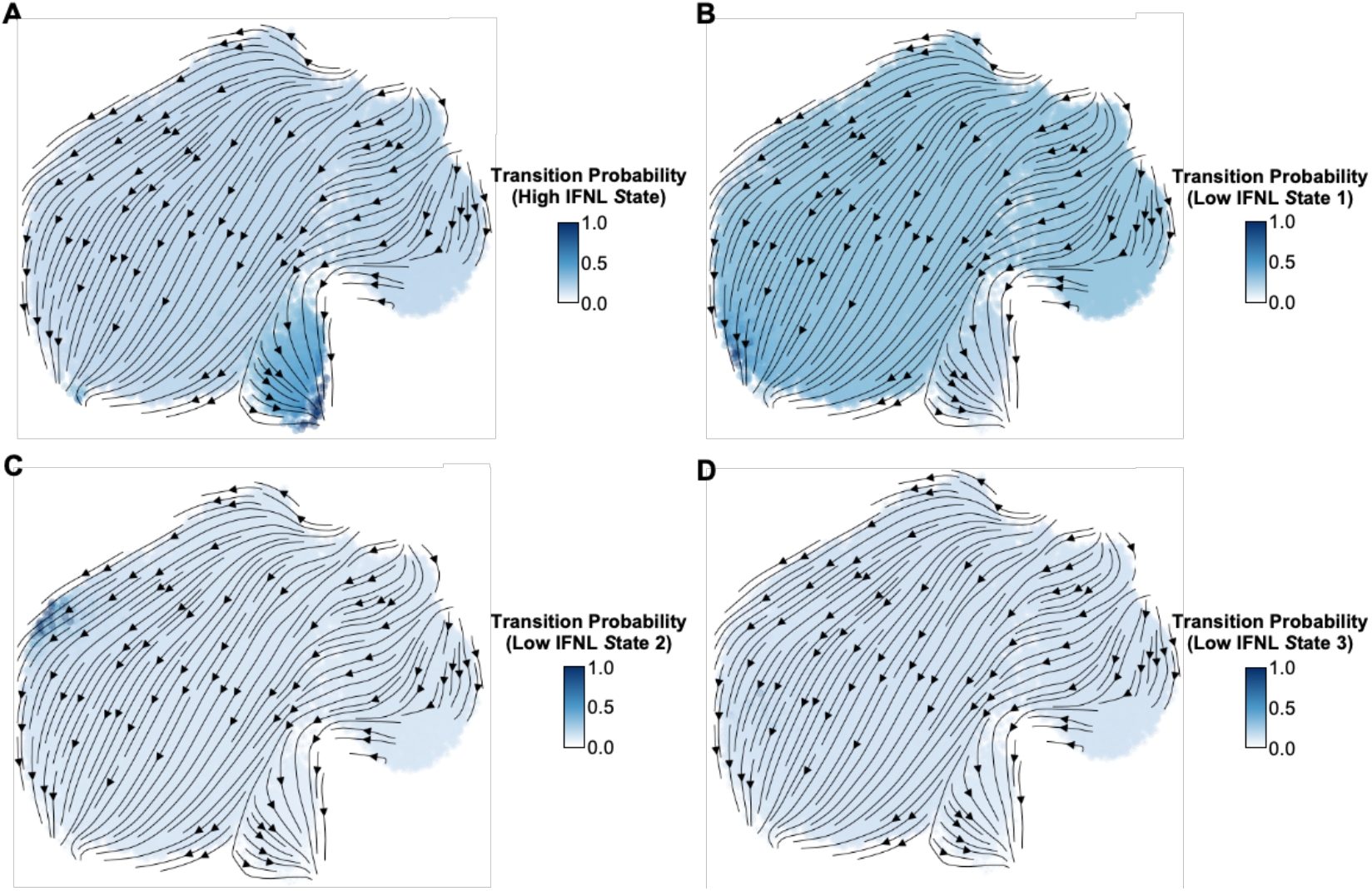
Transition probabilities for all cells transitioning to terminal states. **A)** Maps of the scRNA-seq dataset highlighting transition probabilities for high IFNL state 1. **B)** scRNA-seq data colored by transition probability to low IFNL state 1, **C)** low IFNL state 2, and **D)** low IFNL state 3.

**S3 Fig.**
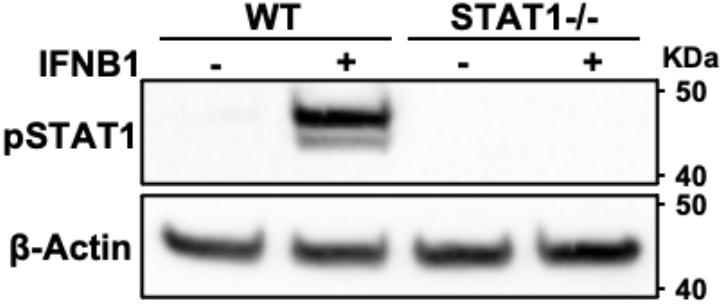
Validation of STAT1-/-A549s. Levels of pSTAT1 in WT or STAT1-/-cells treated with IFNB1 (100 ng/mL) for 16 hrs.

**S4 Fig.**
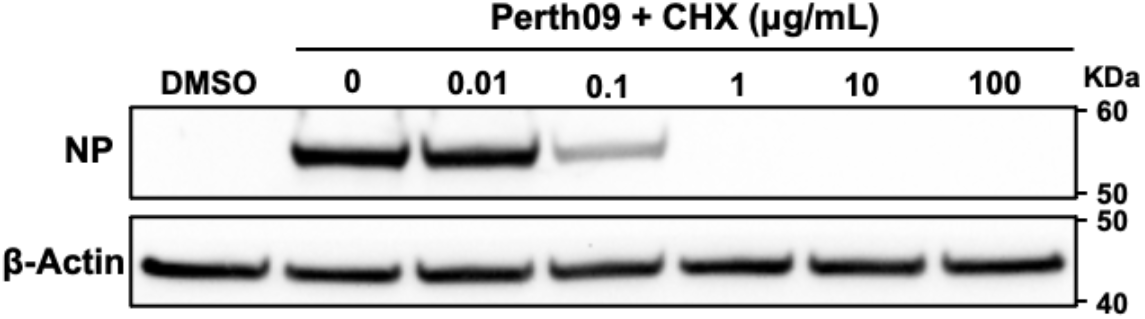
Determining dose of cycloheximide inhibiting translation in A549s. Levels of viral nucleoprotein (NP) in Perth09-infected A549 cells and subsequently treated with difference concentrations of cycloheximide (CHX) for 16 hrs.

**S5 Fig.**
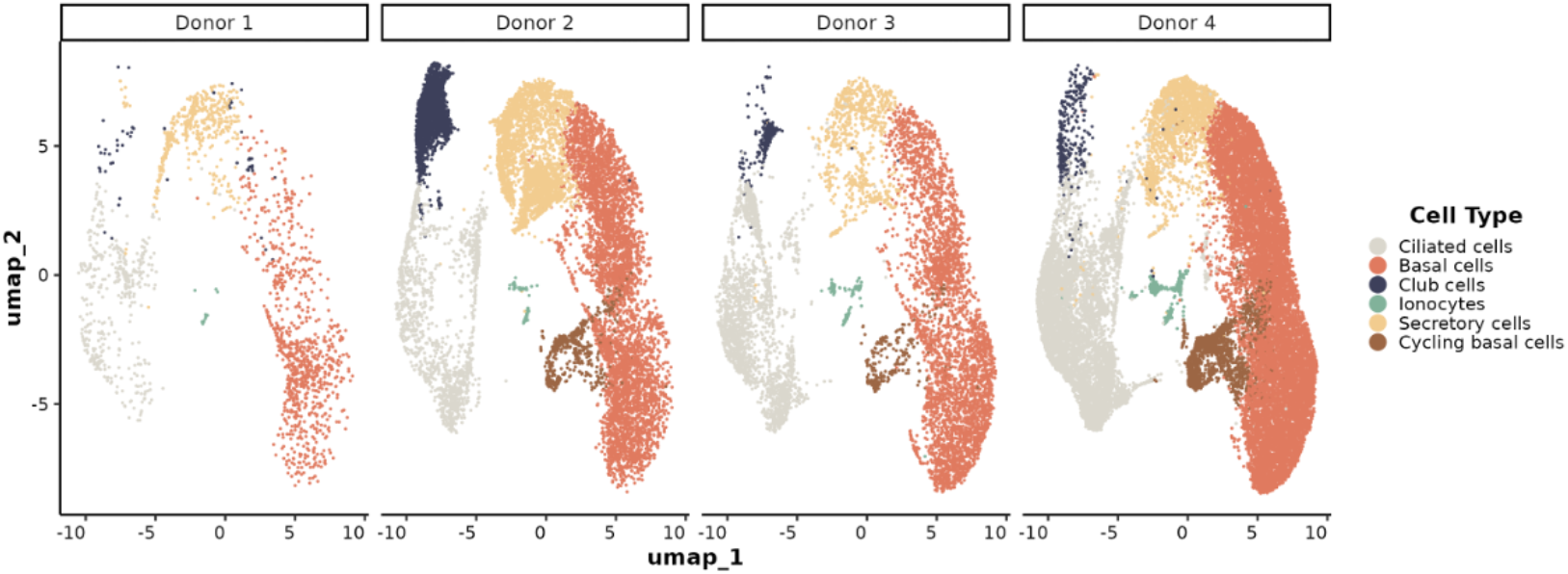
HBECs dataset obtained from four different donors. UMAP representation of scRNA-seq libraries from HBECs split by donor and colored based on cell types present in the population.

**S6 Fig.**
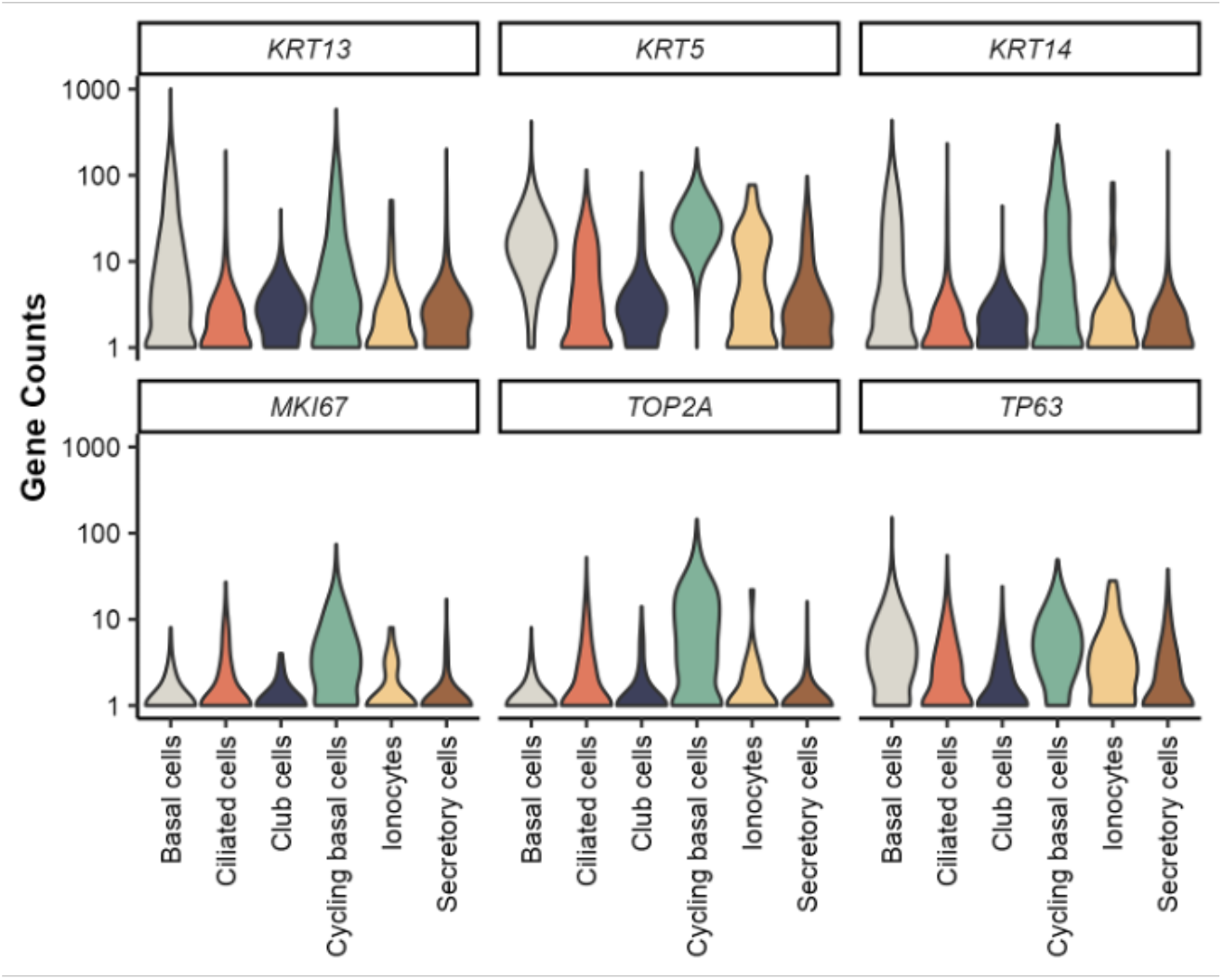
Identification of cycling basal cells in HBECs based on transcriptional profile. Quantification of expression levels of genes markers for basal cells (*KRT13, KRT5, KRT14* and *TP63*) and proliferation (*MKI76* and *TOP2A*) across the different cell type population in scRNA-seq library from uninfected primary human bronchial epithelial cells (HBECs).

